# Mapping the spectrotemporal regions influencing perception of French stop consonants in noise

**DOI:** 10.1101/2024.06.25.600732

**Authors:** Géraldine Carranante, Charlotte Cany, Pouria Farri, Maria Giavazzi, Léo Varnet

## Abstract

Understanding how speech sounds are decoded into linguistic units has been a central research challenge over the last century. This study follows a reverse-correlation approach to reveal the acoustic cues listeners use to categorize French stop consonants in noise. Compared to previous methods, this approach ensures an unprecedented level of detail with only minimal theoretical assumptions. Thirty-two participants performed a speech-in-noise discrimination task based on natural /aCa/ utterances, with C = /b/, /d/, /g/, /p/, /t/, or /k/. The trial-by-trial analysis of their confusions enabled us to map the spectrotemporal information they relied on for their decisions. In place-of-articulation contrasts, the results confirmed the critical role of formant consonant-vowel transitions, used by all participants, and, to a lesser extent, vowel-consonant transitions and high-frequency release bursts. Similarly, for voicing contrasts, we validated the prominent role of the voicing bar cue, with some participants also using formant transitions and burst cues. This approach revealed that most listeners use a combination of several cues for each task, with significant variability within the participant group. These insights shed new light on decades-old debates regarding the relative importance of cues for phoneme perception and suggest that research on acoustic cues should not overlook individual variability in speech perception.

## Introduction

This study investigates the acoustic information on which the human auditory system relies to decode a speech signal into phonemes. Bridging the realms of acoustics and linguistics, speech can be seen as a code between specific time-varying spectral patterns, called acoustic cues, on the one hand, and elemental linguistic units, such as phonetic traits, on the other hand. This “acoustics-to-phonetics conversion” forms the first layer of the recognition chain, whose output is then transformed into phonemes, syllables, words, etc., to finally retrieve the original meaning of the utterance. The relationship between the acoustic signal and its decoding into linguistic units has been a key research problem for speech science in the last century. However, as of today, there is still an important open debate on the exact acoustic information extracted and exploited by our perceptual system to recognize and categorize phonemes. In this article, we reveal the spectrotemporal mapping of the acoustic information listeners use to categorize (French) stop consonants in noise by analyzing the confusions produced at the trial-by-trial level. While other methods based on signal reductions have already been used to pursue this goal (see below for a review), this type of approach offers the potential to achieve an unprecedented level of detail in localizing of the acoustic cues. Over the course of a century of scientific investigations on the acoustic cues governing stop consonant recognition, different methodologies have been proposed, yielding convergent yet not entirely congruent insights. Historically, Fletcher ran in 1922 the first perceptual experiments in order to identify acoustic information in the speech signal^1^. He had participants listen to and repeat nonsense speech syllables, either low-pass or high-pass filtered, with various cutoff frequencies. In this way, he sought to identify the frequency bands which contribute the most to intelligibility. Although this initial approach lacked the precision needed to examine individual acoustic cues, Fletcher’s work laid the foundation for subsequent developments in experimental phonetics. Notably, it led to the important conclusion that the speech signal encompasses multiple cues distributed across independently-processed frequency channels and optimally combined^2,3^.

Following Fletcher, other teams measured the intelligibility of nonsense syllables at various signal-to-noise ratio (SNR). In particular, the examination of confusion matrices in syllable-in-noise recognition tasks by Miller & Nicely^4^ revealed that perceptual errors were not randomly distributed. Instead, these errors clustered within confusion groups aligned with primary phonological features identified by phoneticians. For instance, stop consonants in French are organised along two main features: voicing, distinguishing between voiced ([b], [d], [g]) and unvoiced ([p], [t], [k]) stops, and place of articulation, distinguishing between labial ([b], [p]), coronal ([d], [t]) and dorsal ([g], [k]) stops. A careful examination of confusion matrices as a function of SNR led to important findings about the acoustic cues underlying the recognition of these consonants^5,6^. Notably, the dental stop [t] exhibited more robustness to noise compared to its labial and coronal counterpart [p] and [k], attributed to the presence of a high-frequency burst cue at the consonant onset. Singh & Allen^6^ argued that the recognition of stop consonants in noise generally depends on the detection of a single necessary and sufficient cue, resulting in an overall binary error pattern (cue detected vs. cue not detected). Exceptions to this rule-of-thumb included the labial stop [b], which seemed to lack a single noise-robust cue, and instances of poorly-articulated utterances introducing conflicting cues into the recognition process.

In the 1950s, Liberman, Delattre, Cooper, and other researchers at the Haskins Laboratories initiated a series of landmark studies on the acoustic cues of stop consonants leveraging one of the earliest speech synthesizers. Based on the spectrographs of natural speech recordings, it was postulated that two cues could be both necessary and sufficient for the perception of stop consonants: the release burst (high frequency energy at the onset of the consonant) and the consonant-vowel transition, which corresponds to the movement of the formants from the articulation of the consonant to the steady state of the vowel (henceforth CV transition). Two experiments were carried out to test this hypothesis. In the first experiment, simplified synthetic speech stimuli were created, composed of a stable vowel part (two formants F1 and F2 at fixed positions) preceded by a burst at varying frequencies^7^. Identification scores measured on a group of listeners revealed that while the burst frequency influenced the perceived consonant, the relationship was intricate and not straightforward. In some cases the distribution of responses was even found to be bimodal, with the same percept emerging for separate ranges of frequencies. The second experiment dropped the burst and examined the effect of CV formant transitions alone^8^. The speech sounds were again synthesized from two formants (F1 and F2). However, this time, the F2 was designed with various transition types, from a rising F2 to a falling F2 with regularly-spaced intermediate steps. The F1 was designed to show either a large or reduced transition to evoke the perception of voiced or unvoiced consonants, respectively. In general, results on the F2 transition indicated that stop consonants were ordered from [b] to [d] to [g] along the voiced continuum, from rising to falling F2, and from [p] to [t] to [k] along the unvoiced continuum. However, it again seemed impossible to find a direct correspondence between phonemes and F2 onset frequency.This has led the authors to suggest that phonetic decision could be based on the combination of several cues. In an attempt to summarize the results obtained by the early synthetic speech experiments carried out at the Haskins laboratories, Delattre^9^ listed three possible acoustic cues for place perception, and six acoustic cues for voicing perception. Later technical developments of speech resynthesis algorithms allowed for a refinement of the “synthetic speech continuum approach” in terms of naturalness of the stimuli. Cue-trading experiments^10^ and 2-dimensional continuum experiments^11^ have also considered the orthogonal manipulation of a primary and a secondary cue, for example formant transition and release burst for /b/-/d/ categorization, or voice-onset time (VOT) and f0 onset for /d/-/t/ categorization^12,13^.

Two years after Liberman et al.’s pioneering synthetic speech experiment, a different approach was proposed to answer the question of whether the burst or the formant transitions was the most important cue for the perception of stops. Contrary to the speech synthesis approach, this method employed natural speech utterances as stimuli. Segments of the sounds were selectively cut out and presented to the participants either in isolation or combined with different contexts. By varying the portion of the signal that was removed, it became possible to locate the temporal position of the necessary and sufficient cues. Using this “gating” approach on unvoiced stops in CV sequences, Schatz demonstrated that the burst cue does not provide sufficient information to identify [k] in all vowel contexts^14^, confirming Liberman et al.’s^7^ conclusion that both the burst and formant cues are required for correct identification of the sounds. Similar phenomena were observed for other stops and other vocalic contexts^15–18^, although other authors have claimed that the burst may actually be a sufficient cue^19,20^. More generally, an important insight from the gating experiments is that, in some cases, recognition scores remained above chance even when the consonant segment was entirely removed, indicating that listeners can make use of the coarticulation cues information located in adjacent segments^16,21,22^.

Recently, Jont Allen and his research group integrated the filtering, gating and masking approaches into a unified framework termed “three-dimensional deep search”. By combining the recognition scores obtained along the frequency, temporal, and intensity dimensions on individual utterances, they were able to identify necessary and sufficient cues with an unprecedented level of precision^23^. In a first study, they observed that the high-frequency burst energy played a necessary role in the recognition of [t]: intelligibility scores for that consonant rapidly dropped when this acoustic feature was filtered out, masked by noise or gated out^24^. Furthermore, the audibility of this cue was shown to directly predict the robustness of consonant [t] in white or speech-shaped noise. This approach was then generalized to all stop consonants^23^. The burst, characterized by its frequency distribution and delay to the onset of voicing, was found to be necessary and sufficient for the identification of all consonants. However, in the case of [d], [g], [b], and [p], the CV F2 transition cue appeared to be an important cue, necessary to achieve perfect intelligibility. The conclusion that stops are primarily defined by the characteristics of their burst was later confirmed by selectively removing this cue and showing that this manipulation could shift perception from one consonant to the other^25^.

As detailed in the previous paragraphs, the extensive exploration of stop consonant perception using various approaches yielded important findings. In particular, perceptual experiments have revealed that the recognition of specific phonemes relies on the detection of fine time-frequency features, while large portions of the speech signal are not information-bearing. Notably, place-of-articulation cues may include (depending on the specific consonant, vocalic context, and parameters of the experiment): frequency of the release burst, F2 CV transition, F3 CV transition, as well as VC transition cues^9,26^. As for the perceptual cues to voicing, they include the intervocalic interval duration, the duration of the preceding vowel, burst strength and location, *f*_0_ and F1 transitions, and the presence of a voice bar^13,27,28^. Despite the overarching consensus on the existence and importance of specific acoustic cues for consonant perception, disagreement persists regarding the relative weighting and exact roles of these cues in the recognition process. Regarding place perception, for instance, some researchers have argued that F2 is the primary cue, complemented by a secondary burst cue^7,8^: although either the release burst or the CV formant transitions alone are sufficient cues to place^17^, the latter have been shown to dominate place perception^29,30^. On the contrary, other authors have claimed that the burst is the dominant cue: according to their results, this cue is context-independent^19^, used in isolation in [t] and [k]^23,24^ and necessary for the correct perception of all stops^6,23,25^.

These divergences in the results of experimental studies may, in part, be due to methodological limitations. Most of the aforementioned experiments used “reduced speech” stimuli, i.e, speech sounds that have been drastically modified to reduce the number of cues they contain. Whether achieved through filtering, truncation or synthesis, this approach makes it possible for the experimenter to manipulate acoustic cues independently, or to isolate specific cues. However, a notable drawback is the often low quality of the reduced stimuli, because of the limited number of cues or because these cues do not co-vary in the same way as in natural speech. It is therefore possible that the cues identified in these experiments are specific to the type of stimuli used, and that they would not generalize to natural speech comprehension. Furthermore, these techniques make it difficult to study several cues at once, which may also explain why different research teams have converged to different cues. One major step towards a more unified approach has been taken by Jont Allen and his team who combined different forms of reduced speech (filtered speech and truncated speech) with a masking experiment^23^. Nevertheless, combining the results of these three types of experiments into a general picture requires a difficult and mostly qualitative interpretation step.

In the present study, we investigated the question of the cues to stop consonant perception using the Auditory Classification Images (ACI) approach, a recently-developed technique for revealing the acoustic cues involved in auditory categorization tasks. Unlike previous approaches, the ACI is based not on reduced speech, but on natural speech stimuli embedded in a high level of background noise with randomly added bursts of energy (referred to as “bump noise”). The rationale behind this approach is to randomly add bursts of noise onto natural speech recordings and to identify which burst locations influence perception the most. The outcomes are summarized into an ACI, a time-frequency map of the impact of burst on phonetic decision. This map therefore indicates which spectrotemporal regions bear information for the task, i.e., corresponds to an acoustic cue. The ACI approach has already been successfully applied to a /aba/-/ada/ categorization task^31,32^, revealing multiple cues for place-of-articulation perception, including the F2 and F1 CV transitions, the release burst, as well as VC transition cues in the initial [a]. The objective of the present study is to extend this approach to all French stops, in order to produce a comprehensive list of the cues used for place-of-articulation and voicing perception in plosive consonants.

The research hypothesis for this study were preregistered (https://osf.io/fqejt). They can be summarized as follows:

**Hypothesis 1** Using the specific set of noises used during the experiments and the derived ACIs, we can predict the trial-by-trial responses (e.g. /ada/ vs. /aga/) of each participant with an accuracy that is significantly above chance.

**Hypothesis 2** The location of the cues revealed by the ACI should match the predictions from the phonetic literature, described in the previous sections, since both are related to the availability of perceptual information to a voicing or place contrast. We thus expect to find significant perceptual weights on the same spectro-temporal regions.

**Hypothesis 3** In each phonetic contrast considered, the ACIs will be globally similar for all participants. In line with the previous ACI studies on /aba/-/ada/ categorization, we expect that the primary cues will be present in every individual while secondary cues may or may not be used by a given listener.

A fourth hypothesis regarding the performances of a perceptual model of the human auditory system in the task was also preregistered. As this analysis is not central to our main conclusions, it will be detailed in the Supplementary materials.

## Methods

All stimuli and procedures described in this section were preregistered (https://osf.io/fqejt). As described in the supplementary materials, the experiment can be replicated within the fastACI toolbox^33^ under the name `speechACI_Logatome` and all analyses and figures can be reproduced using the script `publ_carranante2024_figs` and the data available as a Zenodo repository (https://doi.org/10.5281/zenodo.11507060).

The same methodology as in Osses et al.^31^ was followed, except that 6 new phonetic contrasts and a single noise type (“bump noise”) were used. The readers are referred to this initial publication for a more detailed description of the protocol and analyses. Some of the data presented in the /aba/-/ada/ condition was already reported and analysed in the initial study^31^.

The study was approved by the Comité d’Ethique de la Recherche Paris-Université, and conducted in accordance with the local legislation and institutional requirements. The participants provided their written informed consent to participate in this study.

### Stimuli

#### Target sounds

The experiment comprises 7 phoneme-categorization conditions corresponding to different pairs of stimuli: /aba/-/ada/ (condition ABDA), /ada/-/aga/ (condition ADGA), /apa/-/ata/ (condition APTA), /ata/-/aka/ (condition AKTA), /ada/-/ata/ (condition ADTA), /aba/-/apa/ (condition ABPA), /aga/-/aka/ (condition AGKA).

We used 6 productions of speech consonant-vowel-consonant (CVC) pseudowords from one single male speaker, taken from the OLLO speech corpus^34^ ([aba]: `S43M_L007_V6_M1_N2_CS0`; [ada]: `S43M_L001_V6_M1_N1_CS0`; [aga]: `S43M_L003_V6_M1_N2_CS0`; [apa]: `S43M_L008_V6_M1_N1_CS0`; [ata]: `S43M_L002_V1_M1_N1_CS0`; [aka]: `S43M_L004_V1_M1_N1`). These speech samples were preprocessed to align the time position of the vowel-consonant transitions, to equalize their energy per syllable, and to have the same total duration. The stored sounds have a duration of 0.86 s, and a sampling frequency of 16000 Hz. The target speech sounds are shown in Figure 1.

**Figure 1.**
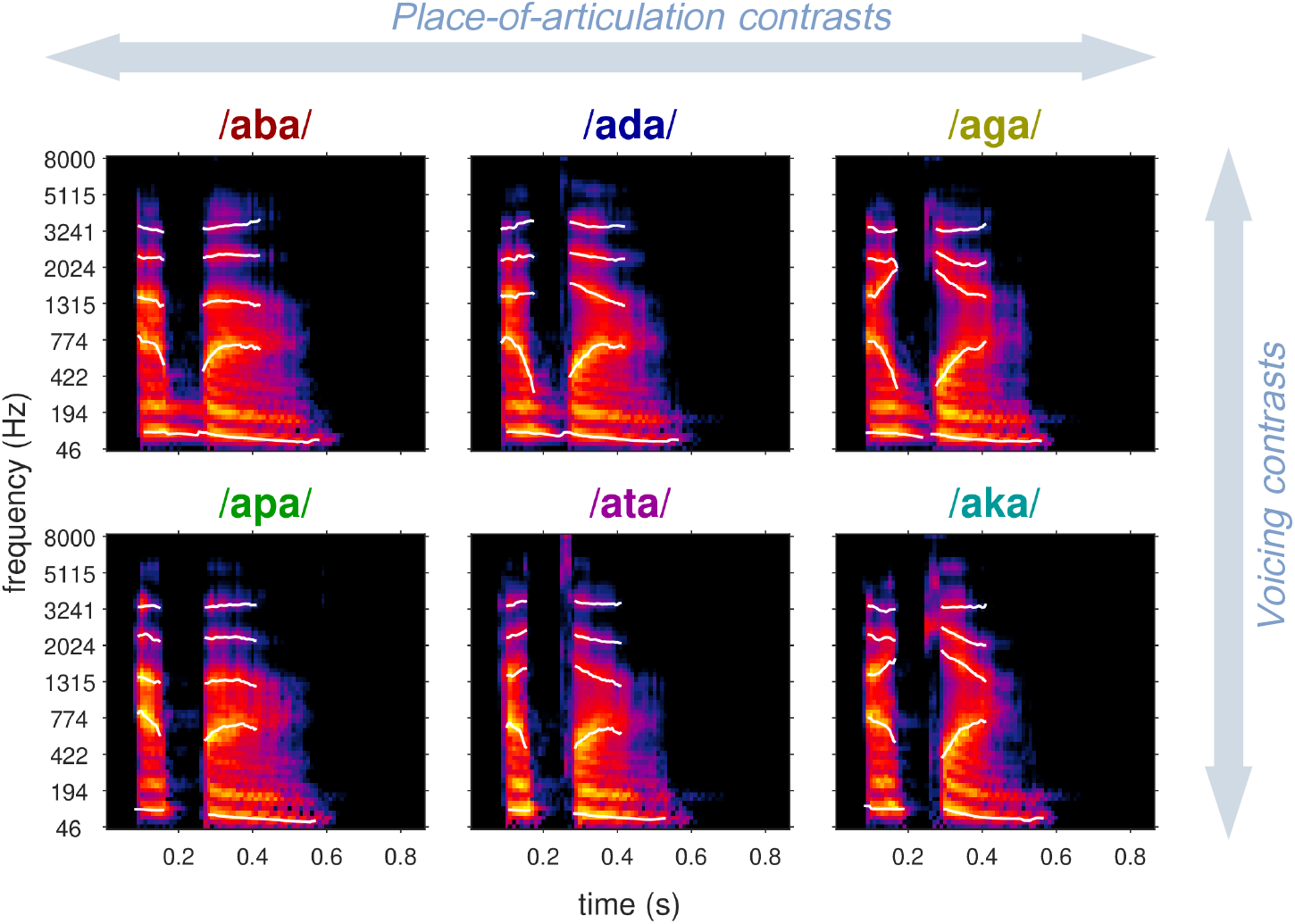
Time-frequency representations of the six targets used in the experiment, organized by distinctive features. The time and frequency resolutions are the same as those used for the analysis. Warmer regions indicate higher amplitudes in a logarithmic scale. The white traces indicate the fundamental frequency ( *f*_0_) and formant trajectories (F1–F4).

#### Background noise

This experiment is based on the “bump noise” condition described in Osses et al^31^. This type of masker has a white-noise-like long-term spectrum, with enhanced temporal fluctuations. The bump noise instances were generated by introducing 30 randomly located Gaussian-shaped bumps into a white noise instance. The bumps had a temporal width of *σ*_*t*_ = 0.02 sec, a spectral width of *σ*_*f*_ = 0.5 ERB, with amplitudes emphasized by a maximum of 10 dB.

For each participant, a new set of 4000 noises was generated, with the same sampling frequency and duration as the target sounds. In each trial, the noise was presented at a level of 65 dB SPL and the target sound was adjusted in level depending on the targeted SNR. The resulting noisy speech sounds were presented diotically via headphones. A small roving in the presentation level between -2.5 and +2.5 dB was applied to discourage the use of level cues.

### Experimental procedure

Each dataset consisted in a total of 4000 observations collected for one participant in one condition. The experiment was divided in 10 blocks of 400 trials each (*≈* 15 minutes per block). Each trial consisted in a one-interval two-alternatives forced choice (“yes/no paradigm”): the participant was presented with one of the two noisy targets and had to categorize it as one target pseudoword or the other by pressing button 1 or 2. A total of 2000 of each of the two targets were presented. The order of the trials was randomized across the whole experiment.

After a correct or incorrect response, the SNR of the next target word was decreased or increased, respectively, following a weighted one-up one-down weighted adaptation rule targeting the 70.7% correct point of the psychometric function. Following each trial, participants received feedback on the correctness of their answer and on their average response bias over the last 100 trials. They were explicitly instructed to minimize their response bias as much as possible.

The actual data collection was preceded by a short training to make sure that the participant correctly understood the task. This “warm-up” session, of a duration left to the participant’s appreciation, was similar to the main experiment except for three additional buttons allowing repeating the noisy speech stimuli or listening to the target sounds in silence. The results of this training was excluded from any further analysis.

### Participants

For each condition, the data from seven participants was collected, yielding a total of 49 datasets. Participants were native French speakers aged 18 or more, with at least one normal hearing ear. This was evaluated by an audiometric test carried during the first test session (Supplementary Figure S1). Only participants with normal-hearing thresholds, i.e., with pure-tone average thresholds equal to or lower (better) than 20 dB HL in their best ear, were retained for the main experiment. Additionally, the results of the first experimental session were used to check that participants are able to perform the task. Participants who did not reach a SNR threshold of -11.5 dB or less in the first session were offered to make a second, then a third try. If they failed again to reach the expected minimum threshold, they were rejected from the analysis. All participants provided written informed consent before participating in the experiment. They were paid on an hourly basis (10 EUR/h).

The data was already collected for 6 participants in the ABDA condition and published in a previous article^31^. Since the primary focus of this investigation was not to compare conditions but rather to examine each condition separately, participants were permitted to perform multiple conditions if desired. Three authors of the study (LV, GC and CC) were also included as participants (S01, S13 and S25, respectively).

### Analysis

#### Performance assessment

Various behavioral metrics were computed from each set of data to assess the overall effect of noise on the listeners’ performance, including correct response rate, SNR threshold, discriminability index and criterion. SNR thresholds were calculated as the median of all reversals in each block of 400 trials, excluding the first four reversals.

#### Auditory Classification Images

Each dataset (one participant in one condition) was processed to obtain one individual ACI.

Two criteria were used to select the trials included in the ACI calculation. In each experimental block of 400 trials, the segment corresponding to the initial convergence of the psychophysical staircase was excluded by rejecting the trials up to the fourth reversal. Subsequently, a number of trials with the most extreme SNRs were removed to balance the occurrence of the two alternative responses in the dataset. These criteria were implemented in Osses et al.^31^ to improve the statistical power of the method. They led to the rejection of *∼* 10% of the trials. The remaining participant’s responses were then processed to obtain an individual ACI.

The time-frequency representation of the noise presented in trial *i* was obtained using a Gammatone auditory filterbank and denoted 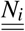. The corresponding response of the listener (“target 1” or “target 2”) is denoted *r*_*i*_. The ACI is a time-frequency matrix of “decision weights” used by the listener to identify the presented phoneme. This matrix of weights, denoted 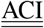, was estimated through a Generalized Linear Model, relating the exact acoustic content of the noise in a given trial to the corresponding response of the participant, according to Equation 1:

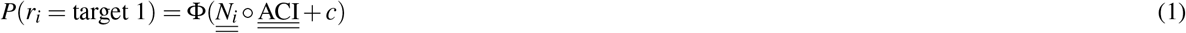

where ○ correspond to the element-wise matrix product, and *c* is an additional model weight representing the participant’s general bias on the considered dataset (by construction, this weight is close to zero). Following the method outlined by Osses et al.^31^, this statistical model was adjusted with a lasso regularization on a Gaussian-pyramid basis. This fitting approach is implemented in the fastACI toolbox under the name `l1glm`.

The above-described analysis pipeline resulted in an individual ACI for each dataset. The significance of each ACI was assessed based on the accuracy of their predictions. In the case of generalized linear models, accuracy is typically reported in terms of deviance, which intuitively measures the deviation from a perfect fit. Here, we report the “mean deviance” obtained following a 10-fold cross-validation procedure (out-of-sample predictive performance of the fitted statistical model described in Equation 1). This deviance is expressed relative to the mean cross-validated deviance of a “zero-weight ACI”, following the same procedure as Osses et al.^31^ Therefore, any deviance value significantly smaller than zero reflects predictions significantly better than chance. This prediction performance metrics was first obtained for each individual ACI (within-subject deviance), then also in cross-predictions between participants engaged in the same condition (between-subject deviance). For this purpose, the ACI of each participant was used to predict the data of each other participant in the same condition. These between-subject predictions offer a way to quantify the similarity among listening strategies within a condition.

The focus of this study being on the exact composition of the ACIs, we need to assess the significance of each perceptual weight. The lasso regression approach can already be seen as a form of “variable selection” where only the relevant variables are associated with non-zero weights. However, we want to confirm this using a significance level threshold. This was done using a randomization approach: a level of weights corresponding to the null hypothesis of no effect was derived by randomizing the responses of the participants and deriving an ACI corresponding to these random responses. This process was iterated 1000 times to obtain a distribution of weights under the null hypothesis. Then the amplitude of the measured weights was compared to this null distribution to determine if they had less than 5% of chance to occur by chance.

Given the small sample size in each condition, no statistical test was performed on the ACIs at the group level. This approach focusing on individual results is in line with most of the reverse correlation literature^35^, and with the experimental phonetics literature^36^.

## Results

In this experiment, six recordings from one single speaker were used in seven phoneme-categorization conditions corresponding to different pairs of stimuli: /aba/-/ada/, /ada/-/aga/, /apa/-/ata/, /ata/-/aka/, /ada/-/ata/, /aba/-/apa/, /aga/-/aka/. Each condition was completed by 7 participants, resulting in 49 sets of behavioral data.

The phonemic contrasts were tested using an adaptive procedure targeting the 70.7% point of the psychometric function. As a result, the average SNR was not the same across the different conditions, reflecting the fact that some discrimination tasks were more difficult than others. Participants performed particularly well in the [g]-[k] contrast (mean SNR = -17.1 dB) and the [d]-[t] contrast (mean SNR = -15.0 dB) while the other conditions resulted in SNRs between -12.9 dB and -13.9 dB (see Supplementary Figure S2). An analysis of variance carried on the SNR thresholds per experimental block confirmed that there was no strong learning effect occurring over the course of the experiment (see Supplementary Analysis). In addition, the performances of the participants in the different conditions were characterized in terms of sensitivity and criterion as a function of SNR. These preregistered analyses are reported in the Supplementary Materials.

One ACI was obtained for each participant and each condition. Each of the 49 individual ACI therefore corresponded to a GLM adjusted to a separate dataset (see Supplementary Figure S3). Figure 2 presents the averaged ACIs per condition, with the formant and f0 trajectories superimposed in order to facilitate interpretation.

**Figure 2.**
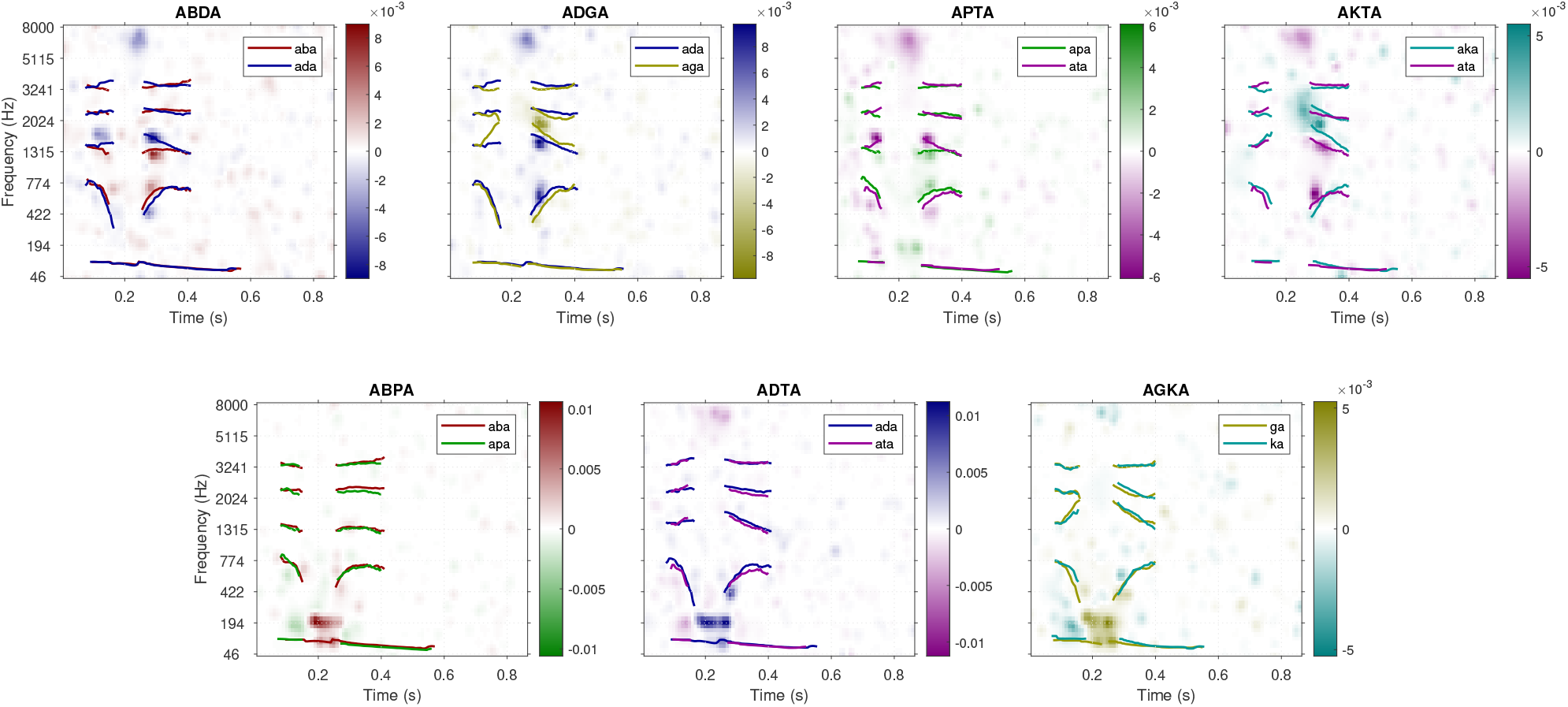
Mean ACIs for all conditions. Top line: place of articulation contrasts. Bottom line: voicing contrasts. The colored lines correspond to the formant and *f*_0_ trajectories for the two targets.

For each individual ACI, the quality of the fit was measured using the cross-validated within-subject deviance metrics (depicted as blue dots in Figure 3) expressed relative to a baseline corresponding to a “null” ACI where all weights are set to zero (dotted line). Remarkably, only two out of the 49 ACIs failed to produce predictions significantly better than chance. Additionally, cross-validated deviance was computed across participants (between-subject deviance) to assess the similarity between individual ACIs within each condition. If two participants use similar listening strategies, the data from one participant should be predicted equally well using either their own ACI or the ACI of the other participant^31,37^. On the contrary, any significant gap between the within-subject and between-suject accuracy indicates that the weighting patterns within the considered ACIs are substantially different. The red dots in each panel of Figure 3 represent how well the data from this participant is predicted by the other participants’ ACIs, on average. This analysis demonstrates that a majority of participants (33 out of the 49 datasets), obtained a between-subject deviance significantly higher (i.e. worse) than their within-subject deviance. Therefore, these participants use a listening strategy that is significantly dissimilar to the other members of the group.

**Figure 3.**
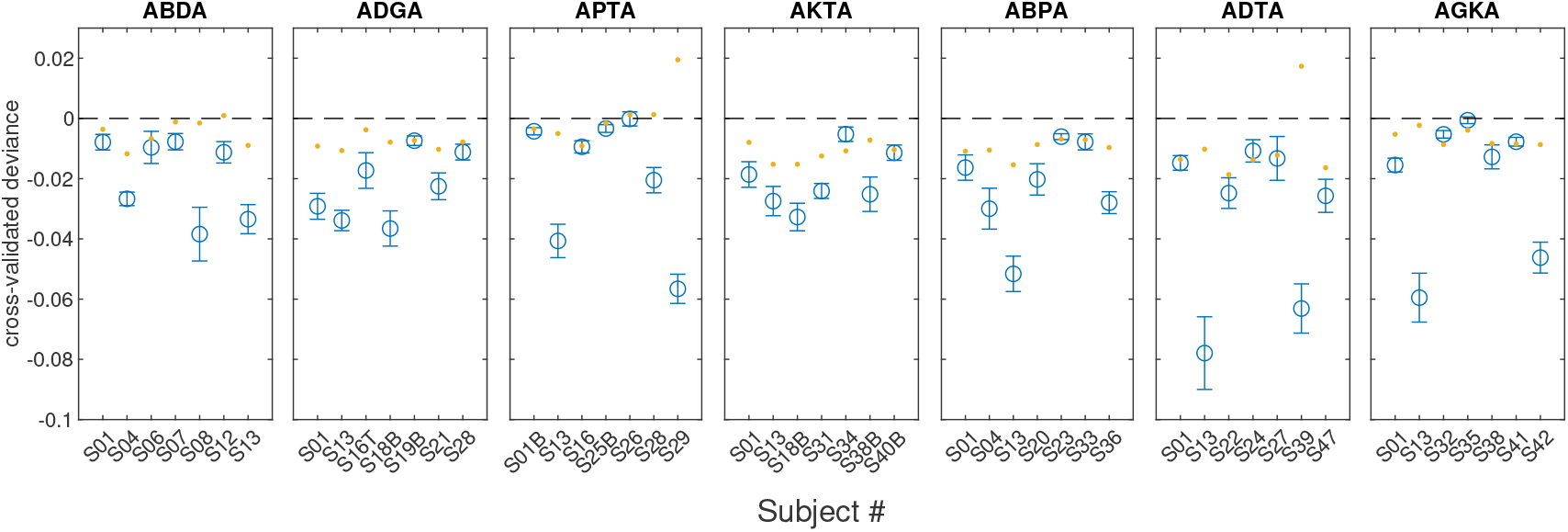
Cross-validated within-subject deviance per trial, for each individual ACI (blue). Error bars represent the variability across cross-validation folds (*±* 1.64 SEM). The deviance is expressed relative to a baseline corresponding to the prediction accuracy of a zero-weight ACI (black dotted line). Therefore, negative deviance values indicate an improvement in prediction accuracy, compared to chance. The red dots correspond to the mean between-subject deviance. This value is obtained by predicting each participant’s data using the ACIs of the 6 other participants in the same condition.

Visual inspection revealed that most of the ACIs corresponding to place-of-articulation contrasts showed a clear pattern of positive and negative weights on the F2 onset, organized vertically – indicating a frequency discrimination. This was confirmed quantitatively as significant weights were found in the region of the F2 onset for all 28 place-of-articulation ACIs. Additionally, weaker weights were found on the F1 onset and on the location of the high-frequency release burst for the coronal stop (> 5 kHz) and the dorsal stop (*∼*2.5 kHz). Furthermore, the labial vs. coronal contrasts appeared to involve some CV transition cues on the initial [a]. The voicing ACIs revealed a simpler pattern, with positive weights concentrated mainly in the region of the f0 during the intervocalic interval, and preceded by negative weights. Some very weak weights were also found on the F1 onset and on the release burst for /t/ and /k/. The significance of the weights was assessed at the individual level. In Table 1, we report the proportion of ACIs displaying significant weights in a time-frequency regions of interest (ROI) centered on the position of each potential cue. For the sake of simplicity, all ACIs were considered in this analysis, including the two that were not significant. The general pattern for each feature remains largely unchanged when considering only the significant ACIs.

**Table 1.** Overview of the cues found in individual ACIs. Number of ACIs for which a given cue was found significant, in each condition.

|  | place-of-articulation contrasts |  |  |  | voicing contrasts |  |  |
| --- | --- | --- | --- | --- | --- | --- | --- |
|  | ABDA | ADGA | APTA | AKTA | ABPA | ADTA | AGKA |
| CV F2 transition | 7/7 | 7/7 | 7/7 | 7/7 | 0/7 | 1/7 | 0/7 |
| CV F1 transition | 3/7 | 6/7 | 2/7 | 3/7 | 4/7 | 7/7 | 4/7 |
| burst | 4/7 | 5/7 | 4/7 | 6/7 | 1/7 | 4/7 | 1/7 |
| voicing bar | 0/7 | 1/7 | 2/7 | 0/7 | 7/7 | 7/7 | 6/7 |
| VC F2 transition | 4/7 | 0/7 | 3/7 | 0/7 | 0/7 | 1/7 | 0/7 |
| VC F1 transition | 1/7 | 0/7 | 3/7 | 0/7 | 3/7 | 2/7 | 2/7 |

## Discussion

The goal of this study is to identify the specific acoustic cues on which individuals rely during phoneme discrimination tasks, with a focus on French stop consonants. By analyzing speech-in-noise recognition data through a reverse-correlation approach, we aim, first, to independently corroborate well-established cues already identified by the scientific community with a method based on natural speech rather than reduced/synthesized speech stimuli. Secondly, we seek to emphasize certain aspects of our findings that could contribute to elucidating and reconciling disagreements within the field.

All participants were able to perform the task at a very low SNR (mean SNR across all participants and conditions: -14.1 dB), with some variability between the easiest discrimination task (AGKA, mean SNR = -17.1 dB) and the most difficult ones (AKTA and ABPA, mean SNR = -12.9 dB). Although the experiment was quite long, no significant improvement in performance was observed over the course of the experiment, allowing us to pull all 4000 trials altogether for the ACI analysis.

For each dataset, corresponding to one participant in one condition, an ACI was derived by fitting a GLM on the trial-by-trial association between a particular noise instance and the corresponding behavioral response of the participant (e.g. /ada/ vs. /aga/). The significance of each ACI can be measured through the prediction accuracy of the statistical model^31^. Prediction accuracy was significantly above chance in 47 out of the 49 ACIs (blue circles in Figure 3), confirming that the model successfully captured some aspects of the listeners’ strategy (preregistered hypothesis 1). As the model gives a reliable account of the participants’ pattern of behavior in the task, it is therefore legitimate to explore the model weights as an insight of the participant’s perceptual strategy.

It is important to note, however, that not all individual ACIs exhibit identical weight patterns. In order to assess the homogeneity of listening strategies (preregistered hypothesis 3), we measured the between-subject accuracy between individual ACIs within each contrast. This analysis demonstrates that a majority of participants (33 out of the 49 ACIs) used a listening strategy that is significantly dissimilar to the other members of their group. For example, S39 appears to rely mostly on high frequency information (*∼* 8 kHz) contrary to other participants in the APTA condition (Supplementary Figure S3).

This variability cannot be explained by obvious differences in hearing threshold or linguistic background as we have selected a homogeneous group of normal-hearing native french speakers. Further study is needed to investigate whether this variability may arise from more subtle differences in hearing thresholds or second languages. Inter-individual variability on perceptual cues in phonological perception has already been documented^38,39^. For example, it has been shown that there is an individual variability in the reliance on the VOT and f0 onset cues for voicing perception in English^13,40^ and for stop perception in Korean^41^. In the next section, we describe the general shape of the ACIs in each contrast. The implications of the interindividual variability for the conclusions will be examined in the rest of the discussion.

### Acoustic cues of French stop consonants

For place-of-articulation contrasts (ABDA, ADGA, APTA, AKTA, top panels of Figure 2), the results are in general agreement with those obtained in prior ACI studies focusing on /aba/-/ada/^31,32,42^ and /alda/-/alga/-/arda/-/arga/^37,43,44^ categorization tasks. The ACIs show a clear pattern of positive and negative weights organized vertically and matching the location of the CV F2 transition for the corresponding targets (/aba/ and /apa/ at *∼* 1300 Hz, /ada/ and /ata/ at *∼* 1700 Hz, /aga/ and /aka/ at *∼* 2000 Hz). The organisation of these weights indicates that when noise energy is present at the time-frequency location of one of the two targets’ CV F2 transition, it influences perception towards that target. This is a marker of the role of the CV F2 transition for place-of-articulation contrast perception, as already shown by the literature on stop consonants using synthetic speech^7,8,45^. In some, but not all, place-of-articulation ACIs, a similar pattern of weights is visible around the CV F1 onset. The role of the CV F1 transitions in stop consonant perception has been noted in the literature, albeit less frequently^45–47^. In our study, this information was used by half of our participants (14 out of 28), therefore confirming that this cue may play a role in a context of speech-in-noise comprehension. As suggested by Ohde and Stevens, it could be the case that the perceptual property critical for place perception is the relationship between F2 and F1 at syllable onset^46^. Symmetrically, positive and negative weights were found on the initial /a/ segment in the labial vs. coronal contrasts (ABDA and APTA), corresponding to the VC F2 transition (for 7 out of the 14 participants) and VC F1 transition (for 4 participant only). VC cues are generally weighted less heavily than the corresponding CV cues, consistent with the notion that the former play a minor role in place perception^22^. In the coronal vs. dorsal contrasts (ADGA and ATKA) these cues are absent, possibly because VC formant transition frequencies show a larger overlap (and therefore a less robust encoding) for back places of articulation^48^. Finally, most place-of-articulation ACIs also reveal a high-frequency cluster of weights at syllable onset, cueing for the detection of [d] and [t]. This indicates that a majority of participants rely on the detection of a burst of energy at the release of the consonant to identify dental plosives, as highlighted by numerous prior studies^23^. There are still ongoing discussions to arbitrate whether F2-transition cues and burst cues should be considered necessary and sufficient cues^23,45,47^. We go into more detail about this matter in the next section.

For voicing contrasts (ABPA, ADTA, AGKA), 20 out of 21 ACIs show weights in the region of the f0 and the first harmonic (*∼* 100 and 200 Hz, respectively), during the intervocalic interval. This low-frequency region corresponds to the voicing bar, which is characteristic of voiced sounds. It is therefore expected to find it as a cue for identification of the voiced stop consonant ([b], [d] and [g]). Additionally, some of the voicing ACIs display weights organized vertically on the onset of F1, indicating a CV F1 transition cue, and weights in the high frequencies for [t] and [k] correspond to release burst cues. Both of these cues have also already been mentioned by the literature on voiced stop consonants^26,27,49,50^.

As detailed above, the acoustic cues revealed by the ACI method closely align with the phonetic literature. The present study therefore confirms the role of these cues for perception of natural (i.e. non-manipulated) speech recordings in noise. However, some previously noted acoustic cues are absent from our data. This is particularly striking in the case of voicing: while some authors have catalogued no less than 16 cues distinguishing between [b] and [p]^27^, our ACIs for the ABPA contrast revealed only one cue shared by all participants. In particular, the *f*_0_ transitions do not seem to be used by any of 21 listeners engaged in the voicing discrimination task. Similarly, the F3 transition and the voice onset time, highlighted as cueing for place perception in past studies^7,8,45^, do not correspond to clear weights in any of our place-of-articulation ACIs.

Several hypotheses could explain the absence of these cues. First, this might be due to the specific set of recordings used in the experiment, which consisted of one single utterance from a single speaker for each of the six target consonants. However, a crucial feature of the ACI method is that it allows to visualize the cues expected by the participant, even when these cues are not actually present in the stimuli^32,51^. Therefore, we believe that the choice of a limited number of utterances per target does not restrict the generality of our results. Further arguments supporting this point are provided in the Supplementary Materials. Another factor that may limit the generality of our conclusions is the choice of a single [aCa] context and a particular language. For instance, since English voiceless stops in this position are typically aspirated, English-speaking listeners might use burst intensity as a voicing cue more systematically than our French-speaking participants. Further studies will be dedicated to extend these results to other contexts and languages.

Second, the method has a limited time-frequency resolution which may restrict the range of observable cues. For instance, a voicing cue on the f0 CV transition may correspond to a region too small to be detected by the lasso regularization on a Gaussian-pyramid basis. Another situation could be that the cues are too close to each other, or even integrated into a single perceptual property. This has been shown to be the case in voicing perception for the F1 and f0 transitions with the voicing bar^28^ and in place perception for the burst and formant transition^52^.

A third explanation involves the interindividual variability observed in our data. Although we initially hypothesized that the individual ACIs of participants in each phonetic contrast would be generally similar, it turns out that some cues are used by all participants (the F2 onset and the voicing bar for the place-of-articulation contrasts and the voicing contrasts, respectively), while other cues are only found in a subset of participants. Consequently, it is theoretically possible that we have not captured the complete range of cues that are used for these contrasts. This, however, is very unlikely: although the sample size per contrast is small, the phonologically orthogonal experimental design minimizes the risk of missing significant cues because they are expected to appear across several contrasts.

A last potential explanation deserves consideration. Most of the psycholinguistic studies discussed so far focus on phoneme perception in quiet, while the present paper is interested in speech-in-noise perception. Accordingly, the experiment is designed to revealing cues relevant to perception in the presence of a white-noise-like background. Given the very low SNR at which participants performed the task, the ACIs likely represent only the most noise-resistant cues^24,47,53,54^. Interestingly, from this perspective, the number of cues revealed by the place-of-articulation ACIs appears to be surprisingly large, compared to previous studies on phoneme-in-noise perception. For instance, according to Régnier & Allen, recognition of [t] in white noise is based almost exclusively on the audibility of the release burst. Conversely, Alwan et al. conclude that formant transition cues become increasingly important for discrimination between labial and alveolar consonants as SNR degrades. The ACIs measured in the present study indicate that, for place of articulation, the multiplicity of cues involved in the decision is preserved up to very low SNRs.

### Cue weighting

Despite a widespread agreement on the existence of specific acoustic cues in consonant perception, there remains ongoing debates about the effective use and the relative importance of these cues in the recognition process. In the context of place perception, for instance, some researchers argue that F2 is the primary cue, supplemented by a secondary burst cue^7,8^. While acknowledging that both the release burst and formant transitions alone can serve as sufficient cues for place identification^17^, they stress that the formant transitions tend to dominate in place-of-articulation perception^29,30^. On the contrary, other authors contend that the burst is the predominant cue, or even for some phonemes like [t], the only cue used for their recognition^23^. According to their findings, this cue is context-independent and essential for accurate perception of all stop consonants^6,19,23,25^. With respect to the relative importance of F2 and the burst cue in place perception, the ACIs measured in the present study reveal that both cues are present in all contrasts (Figure 2, top line). This seems to indcate that for natural speech perception in noise, perception of place is not conveyed by a single critical acoustic feature.

However, it is essential to understand these disagreements within their methodological context. The debate surrounding the use and dominance of acoustic cues in speech perception is deeply influenced by advancements in research techniques and methodologies. Historically, most experimental methods were aimed at identifying the necessary and sufficient acoustic cues for phoneme recognition^6,36^. Experimentally, a necessary and sufficient cue is defined as an acoustic property whose presence/absence in the signal dramatically impacts the recognition of the sound: when the cue is removed by filtering^1^, truncation^14^, or selectively manipulated^8,25^, phoneme identification is impaired, leading to the sound either being unrecognized or misidentified as a different phoneme^23^. As described in the introduction, these methods have been successfully used to reveal the necessary and sufficient cues in a number of phonemes. However, by design, they are not suited for investigating a situation where many cues contribute to the phonetic decision. Even the cue-trading paradigm, widely used for the estimation of the weighting of separate cues, becomes extremely time-consuming when more than two cues are involved^55^ (not to mention that it requires the cues to be known beforehand, making it unsuitable for exploratory research). Similarly, for the 3-dimensional-deep-search method, the presence of multiple, possibly conflicting, cues usually yields complicated patterns of behavior that require difficult interpretation work. As a consequence, these studies have sought for a description of the phonetic decision process based on a minimal number of primary cues, usually one or two, combined with secondary cues, either optional or contextual^49^. Although this conceptual framework has proved useful, it has also led to deadlocked controversies, such as the F2 vs. burst debate.

In contrast, recent theoretical considerations have argued that, depending on the research questions, it may make sense to talk about primary and contextual cues in a parallel way, as they both constitute sources of information that individuals weigh differently^49^, or even that there is no objective basis for a distinction between these two types of cues^56^. The innovative feature of the ACI method for acoustic cues investigation is to allow a data-driven exploration of the information used by a listener when discriminating natural speech sounds in noise. Critically, it does not rely on the notion of necessary and sufficient cue. Rather, the method returns a spectrotemporal map of the regions where the presence of noise influenced perception in a systematic way. Any information contributing to the decision can theoretically be identified in this way, even though this information is not critically needed for the identification of the phoneme. Furthermore, this approach involves a minimum number of assumptions on the nature and number of cues being sought. In the present study, a majority of the ACIs (40 out of 49) reveal the use of more than a single cue, with some listeners relying on at least 4 cues for a given contrast (e.g. S13 in condition APTA), confirming that the phoneme recognition process is not based on the detection of a single acoustic feature but rather involves the analysis of complex spectrotemporal patterns.

As noted above, one striking aspect of our results is the presence of a significant interindividual variability in the pattern of cues extracted by each listener in a given task. This variability is not entirely random; rather, it exhibits a structured pattern that provides insights into the underlying processes. In particular, the F2 CV transition is used by all 28 listeners engaged in a place categorization task while the burst cue is found only in a smaller subset of 19 individuals (Table 1). Therefore, although the ACI method does not formally distinguish between primary and secondary cues, our results indicate that F2 may be said to be the dominant cue over the burst cue for place perception in white noise because it is more prevalent within the tested participants. Furthermore, the burst cue can hardly be considered a “necessary” cue for this contrast, contrary to the claim of some authors^23^, because some listeners appear to not use this cue at all. Similarly, despite the presence of interindividual differences, the voicing bar cue appears in all individual voicing ACIs, except one the one that did not reach significance. This was not the case for the burst cue or the CV F1 transition.

As a conclusion, we think that the use of ACIs offers an appealing methodological avenue for psycholinguistics: in showing in a fine-grained manner, and with only few theoretical assumptions, the information that a particular listener uses in a given task, the ACI approach may be able to reconcile the historical research that has focused on the identification of necessary and sufficient cues and the growing body of work investigating individual differences in the perception of speech sounds.

## Data Availability

This study -including hypotheses, stimuli and procedures - was preregistered (https://osf.io/fqejt). The datasets generated during and analysed during the current study are available as a Zenodo repository: https://doi.org/10.5281/zenodo.11507060. The experiment and all analyses are replicable using the fastACI MATLAB Toolbox, available on Github : https://github.com/aosses-tue/fastACI. Detailed procedure is available on the Supplementary Materials.

## Acknowledgements

This study was supported by the French National Research agency through the ANR grants “fastACI” (Grant No. ANR-20- CE28-0004) and “FrontCog” (Grant No. ANR-17-EURE-0017).

## Author contributions statement

G.C., M.G. and L.V. conceived the experiment. G.C., L.V., C.C. and P.F. collected the data. L.V. analysed the results and ran the simulations. L.V. and G.C. wrote the manuscript. All authors reviewed the manuscript.

## Supplementary materials

### Audiometric thresholds

Audibility thresholds were measured using pure-tone audiometry at six frequencies (250, 500, 1000, 2000, 4000, and 8000 Hz) and had average thresholds between -7.5 (S43) and 15.8 dB HL (S33) in their best ear, meeting our inclusion criterion of having thresholds of 20 dB HL or better. The obtained hearing thresholds are shown in Fig. S1.

**Figure S1.**
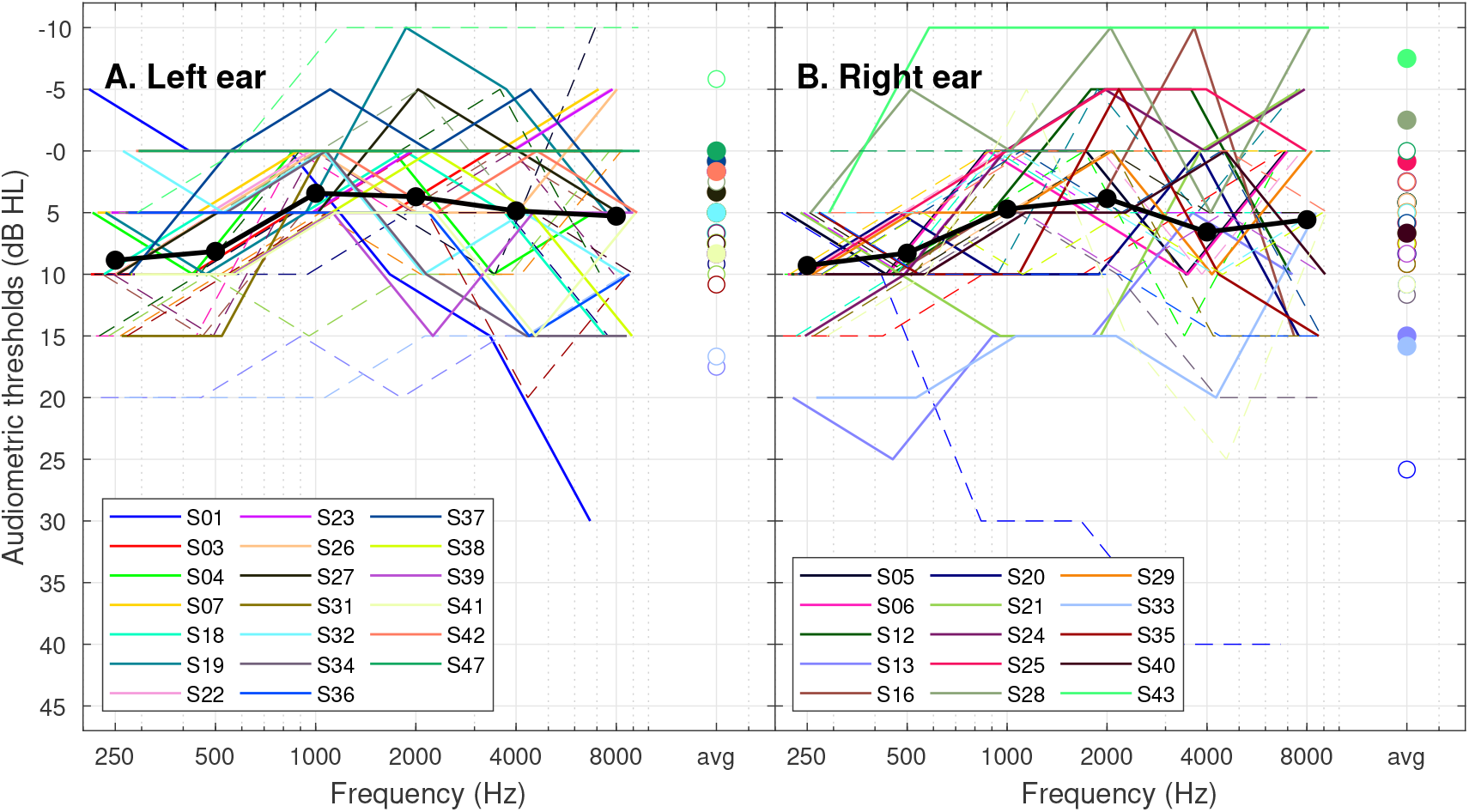
Audiograms for all participants. Left and right ear thresholds are shown in Panels A and B, respectively. The participant’s best-ear thresholds are connected by continuous traces, and the subject ID is indicated in the corresponding panel legend. Average thresholds across participants are indicated by the black traces and the average audiometric threshold for all tested frequencies between 250 and 8000 Hz are indicated by the right-most markers (filled symbols are used when those averages are from the participant’s best ear).

### Behavioral results

The participants’ performances were characterized using several indicators in order to check that they were able to perform the task at a reasonable level (minimum SNR threshold in the 1 block: -11.5 dB), that they were not overly biased in favor of one of the two responses, and that they did not show any strong learning effect over the course of the experiment.

An analysis of variance (ANOVA) was conducted to assess the impact of factors Condition and Session on the dependent variable SNR threshold. Results revealed a significant main effect of Condition (F(6, 420) = 96.29, p < 0.0001), indicating that some constrasts were more difficult to discriminate than others. The main effect of Session was not significant (F(9, 420) = 1.76, p = 0.3705), neither the Condition*Session interaction effect (F(54, 420) = 0.43, p = 0.9998), suggesting small or no learning effect over the course of the experiment.

The performance (discriminability index d’) and bias (criterion) were analyzed for effects of SNR level and condition. For this purpose, the two metrics were represented as histograms, with SNR bins of 1 dB, similar to Varnet & Lorenzi^35^. The plots of d’ and criterion as a function of SNR are shown in figure S2.

Only SNR levels with sufficient data (more than 15 observations for each participant) were considered in the statistical analysis. Two ANOVAs were conducted separately on the dependent variables d’ and criterion. The results demonstrated a significant main effect of Condition (F(6, 165) = 76.4, p < 0.0001) and SNR (F(3, 165) = 66.41, p < 0.0001) on dprime, but no significant interaction effect (F(18, 165) = 1.13, p = 0.33). For criterion, the analysis indicated no significant impact of these factors or their interaction (all p’s>0.3).

### Individual ACIs

Figure S3 shows the individual ACIs for all participants included in the study. For representation purposes, nonsignificant weights are set to zero and all ACIs are normalized in maximum absolute value.

**Figure S2.**
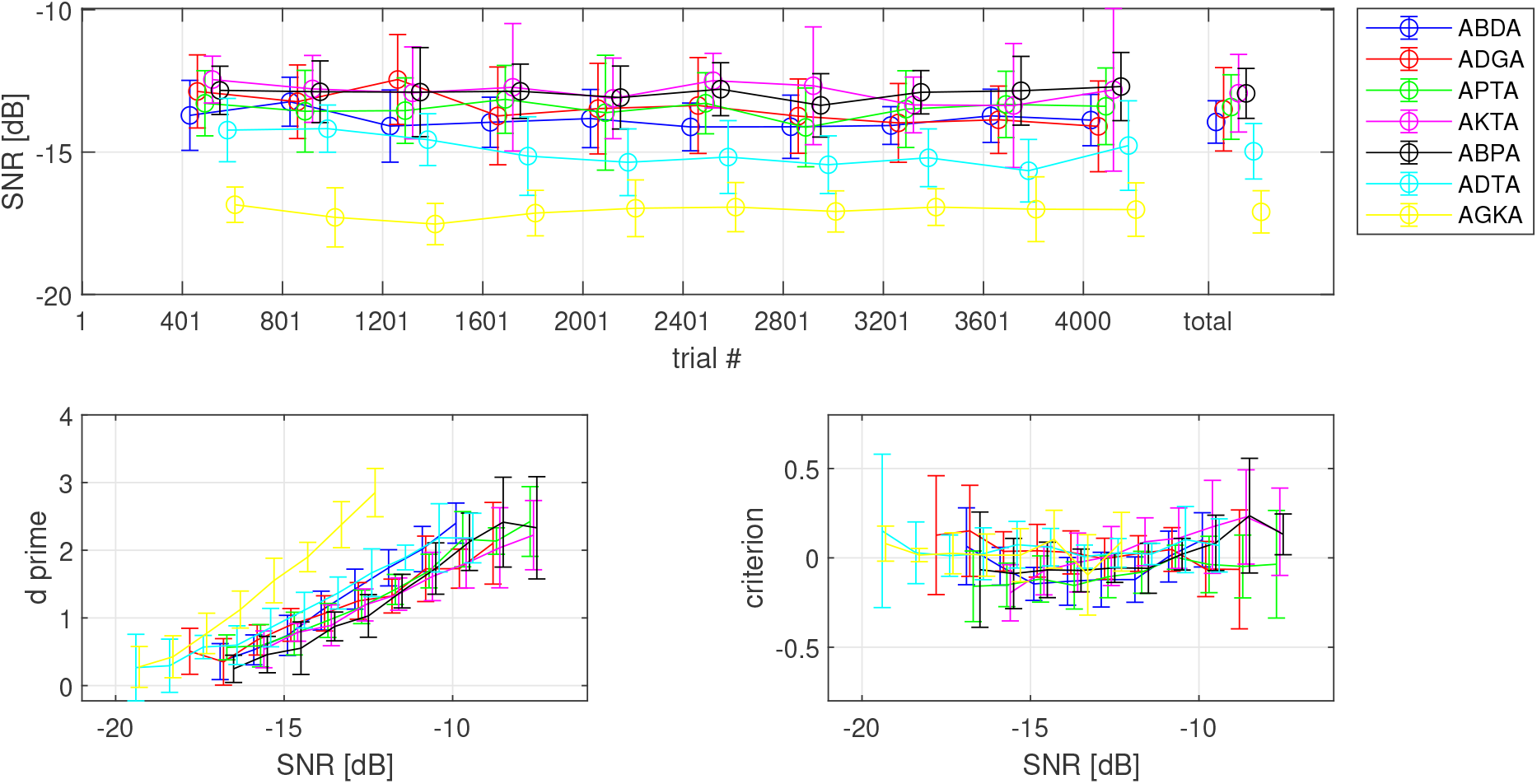
Behavioral performances in the seven conditions. Top panel: SNR threshold over the course of the experiment and overall SNR threshold for the whole experiment. Bottom panels: discriminability index (left) and criterion (right) as a function of SNR. Error bars indicate standard deviation across the seven participants.

### Artificial listener

In addition to the experimental data collection, an auditory model was run on each condition. The artificial listener used in the study was the same as in Osses et al.^31^. This model assesses the internal representations of each sound using signal-driven (bottom-up) information based on a modulation-filter-bank approach. The internal representations are subsequently compared using a simple (top-down) decision back-end based on template matching. The same set of comparison metrics as for “real listeners” was applied to simulated responses from this artificial listener for which ACIs were also derived. The modulation-filter-bank model is implemented in the script `osses2022a.m` and the decision back-end is implemented in the script `aci_detect.m` (option ‘optimal_detector’) in the fastACI toolbox.

Our preregistered hypothesis 4 was that the auditory cues used by humans to perform the categorization tasks would be a subset of the available cues, i.e. of the acoustic differences perceptible by the human auditory system. The use of a perceptual model mimicking the limited resolution of the human ear and using the available information optimally offers a way to test this hypothesis. We also computed the difference between the auditory spectrograms of the targets used in each contrasts.

Figure S4 presents the difference between auditory spectrograms, while the simulated ACIs for each of the seven contrasts are shown in S5.

For place contrasts, the F1 and F2 transitions are clearly visible both in the simulated ACIs and the difference of spectrogram. However, the artificial listener appears to use the temporal dynamics of the formant transitions rather than their onset/offset frequencies, suggesting that its temporal resolution is too high. Other acoustic features were also theoretically perceptible although they were not used by the participants. In particular, the targets present marked differences in the low-freqency region (and to some extent on the F3 and F4) that are not exploited by the participants. For voicing contrasts, the intervocalic region, corresponding to the voicing bar, truly corresponds to acoustic differences between voiced and unvoiced consonants. The artificial listener is able to use this information although it focuses on the onset of the voicing bar – again possibly because of a temporal bias. Similarly, the F1 CV and VC transitions, appear both in the difference of spectrograms and the simulated ACIs. On the contrary, F2, F3 and F4 cues are not used by human listeners whereas the targets exhibit differences that could potentially serve as a basis for discrimination. Finally, it should be noted that the high-frequency release burst cue, that is used by some participants in the place and voicing contrasts, actually corresponds to a very subtle difference in the targets (almost imperceptible in the case of [d]). This cue does not appear in the simulated ACIs, indicating that it was considered too weak by the artificial listener.

In summary, most of the auditory cues used by humans to perform the categorization tasks are a subset of the available cues, i.e. of the acoustic differences perceptible by the human auditory system. This is not entirely true however, as some spectrotemporal regions were actively extracted by the listeners although they did not correspond to any robust, reliable information for the task. This apparent paradox is discussed in the following paragraph.

**Figure S3.**
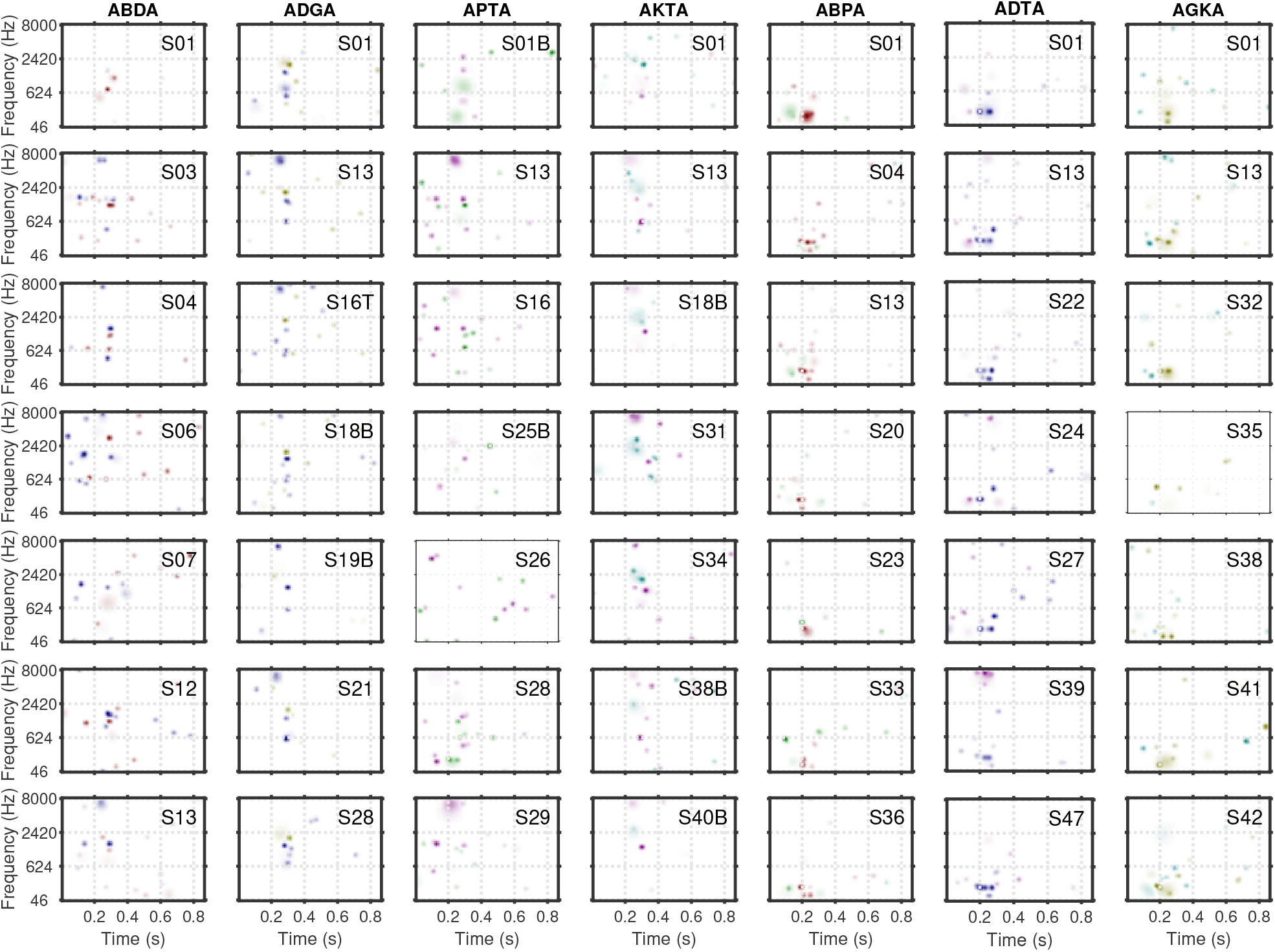
Individual ACIs for all participants in the experiment. Each column corresponds to a different condition. For representation purposes, nonsignificant weights are set to zero and all ACIs are normalized in maximum absolute value. ACIs yielding prediction accuracy significantly above the zero-weight baseline are indicated with bold contours. The color code is the same as in Figure 2.

### Specificity of the results to the target utterances

A potential limitation of the ACI paradigm is its reliance on a single utterance for each target consonant (e.g., one recording of [aba] and one recording of [ada]). This ‘frozen speech’ task is rather unnatural as speech productions typically exhibit a large amount of variability. Moreover, the duration of the experiment may lead participants to rely on target-specific cues rather that general speech cues. It is therefore legitimate to question whether the observed results are specific to the selected set of utterances, or if they reflect broader speech processes. Could it be the case that the cues identified in this study simply stem from incidental acoustic differences between the targets, that the participants would be able to learn to detect? In this section, we provide four arguments countering this claim.

First, a critical feature of the reverse correlation approach lies in its capacity to capture the information represented in the subject’s mind, not only the information available in the stimulus. As Gosselyn & Schyns^51^ stated: “Reverse correlation […] can reveal ‘hidden’ information about a representation in memory when this representation is used to categorize the input”. This has been demonstrated both empirically^51^ and theoretically in the case of linear observers^22,57–59^. As a limit case, the method can work even in the absence of a target, i.e. in noise-alone stimuli, as long as the participant is performing a categorization task on these stimuli^60,61^. In the case of phoneme perception, we have demonstrated that the approach can reveal a phonetic cue expected by the participant even when this cue is not present in the targets^32^. We conducted a [ba]-[da] discrimination task where all targets were preceded by a neutral context /a/, identical for both target. Even so, the measured ACIs revealed VC transition cues in the uninformative initial vowel, demonstrating that the method reflects the listener’s internal expectations rather than the actual acoustic properties of the stimuli^32^. Therefore, the fact that a limited number of targets were used in the present study does not, in theory, restrict the set of potential cues that can be observed. As a matter of fact, the analysis of auditory spectrograms and the simulated data obtained from an artificial listener suggests that the burst cue observed in some ACIs is not actually audible in most of our targets (see previous section).

**Figure S4.**
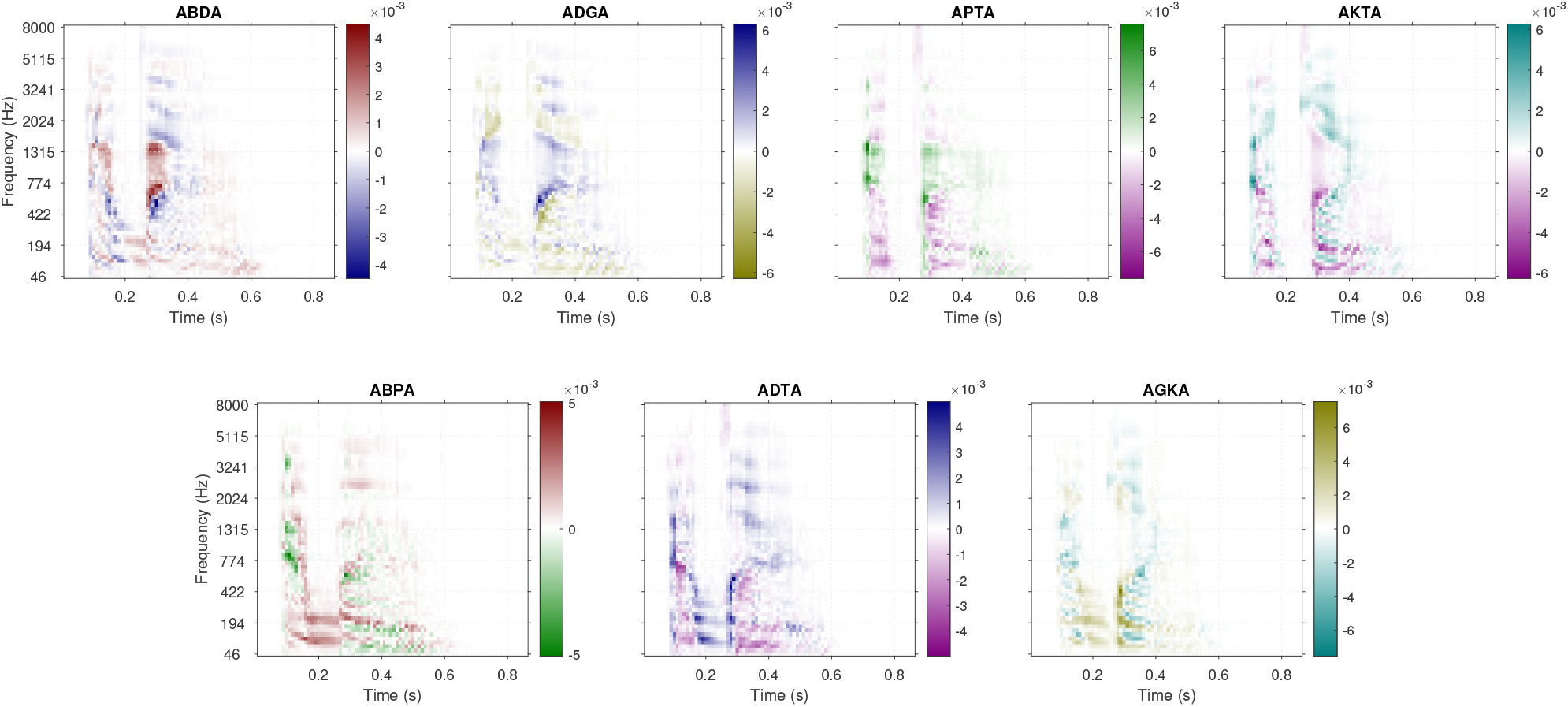
Differences in auditory spectrograms between each pair of target stimuli across all 7 experiments. The color code is the same as in Figure 2.

Second, any learning effect occurring over the course of the experiment, such as the detection of target-specific cues, would likely result in an improvement of the SNR thresholds from one bloc to the next. However, this is not what was observed (see section Behavioral results). Therefore, it is unlikely that the participants learned to detect and use incidental acoustic cues specific to the targets used in the experiment.

Third, according to the simulations and the comparison of auditory spectrograms, the most salient acoustic differences between the stimuli do not necessarily correspond to weights in the ACIs. For instance, the differences in the *f*_0_ trajectory in AKTA are not used by any of the listeners.

Fourth, the measured ACIs are consistent with those obtained in previous studies conducted using different sets of utterances. In particular, CV formant transition cues were observed in all place-of-articulation ACIs measured by our team in past research^11,32,43,62^. Osses and Varnet’s^62^ visual exploration of the ACIs from two listeners in two [aba]-[ada] tasks revealed a generally consistent pattern of weights across different target pairs. A notable exception was the absence of VC transition cues in the results of one experiment. This discrepancy was explained by the fact that the level of the first syllable was relatively low in one pair of targets – and therefore that this target was not audible in the noise masker. Consequently, the participants were effectively performing a [ba]-[da] discrimination, resulting in the absence of any cue in the initial segment.

For these reasons, we do not think that the present experimental paradigm may have prompted participants to use incidental acoustic differences between the utterances as cues for the task. Furthermore, we can reasonably assume that the ACIs provide an comprehensive picture of the cues used for perception of the tested phonemic contrasts in the [a] vocalic contexts at low SNR.

### Replicating the experiment and reproducing the analysis

The fastACI MATLAB toolbox is available on Github at https://github.com/aosses-tue/fastACI. Follow the installation instructions provided in the accompanying file, and initialize the toolbox using the script `startup_fastACI.m`.

For running the experiment, enter the command `fastACI_experiment(‘speechACI_Logatome-abda-S43M’,’S01’,’bumpv1p2_10dB’)` (replace ‘abda’ by the desired condition and ‘S01’ by the subject identifier).

**Figure S5.**
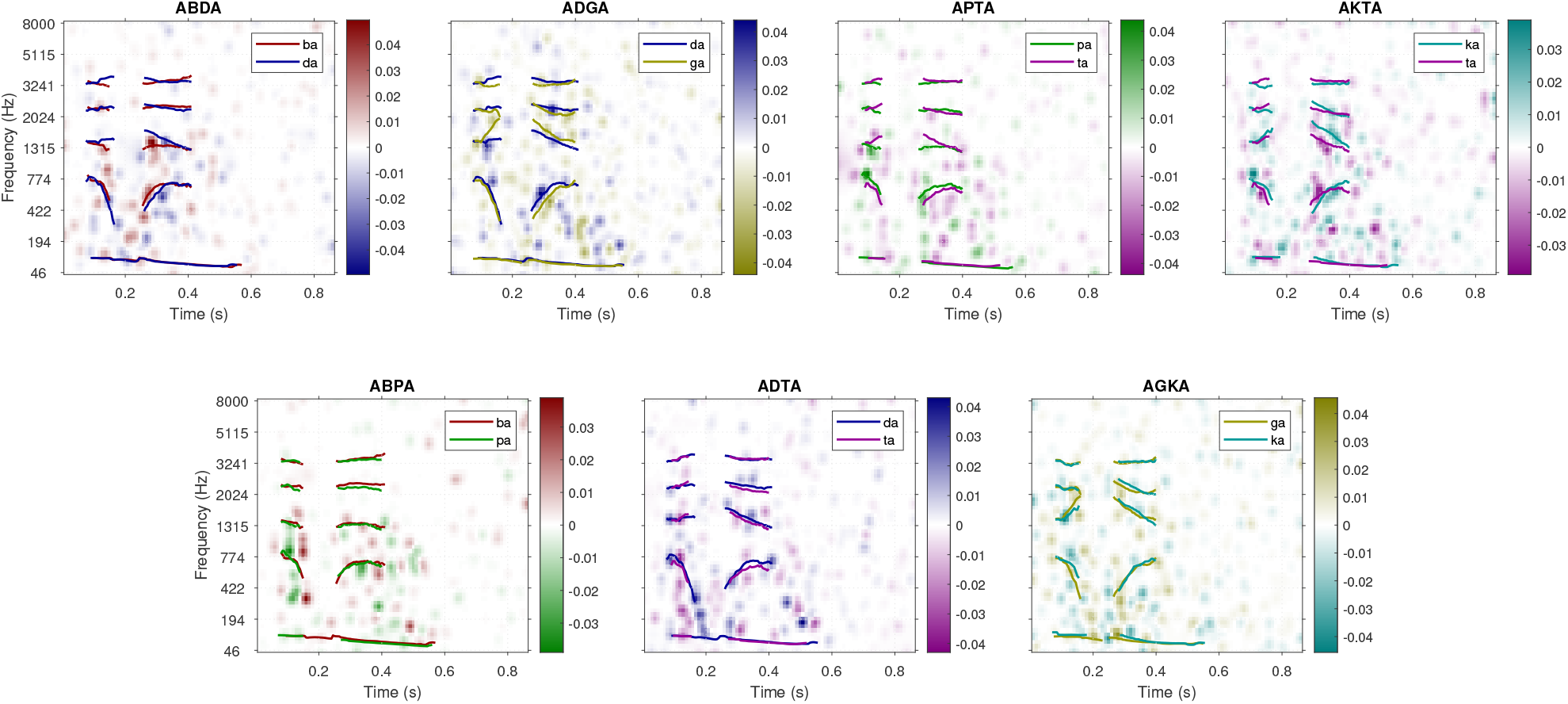
ACIs derived from the simulations using the artificial listener `osses2022a.m`. The color code is the same as in Figure 2.

All figures from the main article and the supplementary materials can be recreated using the fastACI script `publ_carranante2024_figs.` and the name of the figure (e.g. ‘fig1’) as parameter.

If you downloaded the raw and/or post-processed data for this study from Zenodo (https://zenodo.org/records/11507060), you need to specify the extra parameter `‘zenodo’` to indicate the location of the data folder.

